# Novel truncating Desmin mutation Arg150Stop disrupts structural integrity and cellular homeostasis by formation of persistent aggregate-like structure

**DOI:** 10.1101/2025.10.14.682287

**Authors:** Sukanya Mitra, Trisha Ghosh, Arun Kumar Mishra, Shamita Sanga, Mahua Maulik, Moulinath Acharya

## Abstract

Desminopathies are a heterogeneous group of myofibrillar myopathies defined by the presence of desmin-positive aggregates that compromise cytoskeletal integrity in skeletal and cardiac muscle. Although desmin knockout models and several truncating mutations typically result in a functional null phenotype without inclusion body formation, the molecular consequences of specific stop-gain variants remain poorly understood.

In this study, we investigated the pathogenic mechanism of a novel DES nonsense mutation, NM_001927.4:c.448C>T; p.(Arg150Stop), previously identified in an Indian patient with congenital myopathy. This premature stop codon lies within the Linker 1A domain and is predicted to generate a truncated protein lacking the C-terminal tail. To delineate its functional consequences, we used two complementary experimental approaches: transient overexpression of the R150X mutant in skeletal and cardiac myocytes, and a CRISPR/Cas9-engineered homozygous R150X cardiomyocyte line (Des-R150X-CRISPR).

Both models consistently revealed the formation of persistent aggregate-like structures, in striking contrast to desmin knockout systems that do not generate inclusions. These aggregate-like structures disrupted actin filament organization, impaired filament bundling, and induced organelle mislocalization. Biochemical analysis indicated that the aggregates were resistant to proteasomal degradation, yet they were partially cleared by autophagy, underscoring a role for protein quality control pathways in modulating disease severity. Importantly, the Des-R150X-CRISPR line demonstrated aggregate-driven pathology at endogenous levels, confirming that this mutation acts through a toxic gain-of-function mechanism rather than simple loss of desmin function.

Our findings establish the Arg150Stop variant as a mechanistically distinct truncating mutation that generates aggregation-prone protein rather than a null state. By reproducing hallmark features of desminopathy in a physiologically relevant human cell models, this work not only broadens the known pathogenic spectrum of DES variants but also highlights aggregate formation as a central driver of cellular dysfunction and a promising therapeutic target in desminopathies.

## Introduction

Desmin, a 53-kDa type III intermediate filament (IF) protein, is the major muscle-specific IF expressed in cardiac, skeletal, and smooth muscle, serving as a central architectural and functional element (1,2). IFs such as desmin provide mechanical resilience and structural integrity through a robust cytoplasmic network (1). Its exclusive expression in muscle underlines why desmin dysfunction profoundly disrupts muscle physiology (3). Structurally, desmin follows the tripartite IF organization with a conserved α-helical coiled-coil “rod” flanked by variable head and tail domains (4). The N-terminal head is essential for filament assembly, while the C-terminal tail governs network organization and protein interactions (5). Within muscle fibers, desmin monomers dimerize and assemble into a three-dimensional filamentous network around Z-discs. This scaffold interconnects adjacent myofibrils and anchors the contractile apparatus to the sarcolemma (costameres), extracellular matrix, intercalated discs, and organelles such as nuclei and mitochondria (6,7). Thus, desmin ensures mechanical stability, preserves cell architecture, regulates organelle positioning, and contributes to protein quality control (1,6).

Mutations in *DES* gene cause desminopathies, clinically heterogenous myofibrillar myopathies that are often accompanied by dilated cardiomyopathy (3). It is characterized by skeletal muscle weakness, which is slowly progressive and predominant in the distal rather than proximal muscles and in lower limb than in upper limb (8). A defining pathological hallmark of desminopathies is the abnormal accumulation of desmin within muscle fibers, forming amorphous eosinophilic deposits that appear as granular or granulofilamentous material upon electron microscopic examination (8). Biochemical studies show that aggregated desmin adopts amyloid-like fibrils that are cytotoxic to muscle cells and severely disrupt sarcomeric organization (9). Cellular protein quality control relies mainly on the ubiquitin–proteasome system (UPS), which degrades soluble misfolded proteins, and the autophagy–lysosome pathway, which clears defective organelles and larger protein aggregates. Although both pathways are essential for clearance of aggregation prone proteins, desmin-containing aggregates often overwhelms these pathways leading to impairment of proteasomal activity (10) and resistance to autophagic clearance (7), leading to their persistence and contributing to progressive myofibrillar degeneration.

Most pathogenic DES mutations cluster within the highly conserved α-helical rod domain, which is essential for filament assembly and cytoskeletal integrity. Single-point mutations in this region frequently disrupt polymerization, leading to intermediate filament collapse and cytoplasmic aggregates (11–15). The majority of truncating mutations are located in the C-terminal region and trigger nonsense-mediated mRNA decay (NMD), resulting in a complete loss of desmin protein and a classical loss-of-function phenotype (6,16–18). Consistently, desmin-null mice and cell models also exhibit a complete absence of desmin, confirming that loss of the protein alone disrupts filament assembly and cytoskeletal integrity (19,20). However, these desmin-null models fail to fully recapitulate the disease phenotype associated with aggregation. There are also reports of truncating mutations in the linker 1A region and 1B domain, such as p.K144X (16,21) and p.L200fsX20 (16), but their effects on aggregation and the molecular mechanisms driving disease pathogenesis have not been characterized, leaving a gap in understanding the cellular consequences of such truncating variants. Along these lines, the molecular and cellular effects of the novel R150X truncating mutation, previously reported from our lab (22) and located within the linker L1A—a critical segment for filament flexibility and stability—remain unexplored.

To address this, we employed two complementary experimental systems: transient overexpression of R150X desmin in skeletal myocytes and cardiomyocytes, and a heterozygous Des-R150X CRISPR cardiomyocyte model. In the R150X mutant exhibited perinuclear desmin-positive structures, disrupted cytoskeletal architecture, and mislocalized organelles, which were partially resolved upon autophagy activation. Interestingly, patient muscle tissue harbouring R150X showed no detectable desmin (23), presumably due to efficient NMD or proteasomal/autophagic clearance, whereas our overexpression model revealed pronounced perinuclear desmin-positive inclusions. Similarly, the heterozygous Des-R150X CRISPR model also exhibited aggregate-like structures. Taken together, these observations suggest that R150X behaves as a conditional gain-of-function mutation, distinct from other truncating DES variants that undergo NMD, justifying its detailed molecular characterization and highlighting the importance of cellular context in interpreting desminopathy mechanisms.

## Materials and methods

### Cell Culture and Cell lines

Primary human skeletal muscle cells (HSkMC; ScienCell Research Laboratories™) were cultured in Skeletal Muscle Cell Medium (SkMCM) supplemented with 5% fetal bovine serum (FBS; ScienCell), 1% Skeletal Muscle Cell Growth Supplement (SkMCGS), and 1% penicillin-streptomycin (P/S) antibiotic solution. Human cardiomyocyte cell line AC16 (ATCC®) was maintained in DMEM/F-12 medium supplemented with 12.5% FBS and 1% Antibiotic-Antimycotic Solution (100X). Human Embryonic Kidney (HEK293) cells (ATCC®) were cultured in DMEM supplemented with 10% FBS, Gibco and 1% Antibiotic-Antimycotic Solution (100X). All cells were cultured at 37 °C in a humidified atmosphere containing 5% CO_2_.

### Gene Cloning and Recombinant Vector Construction

The coding sequences for full-length wild-type desmin (pCI-HA-WT) and the mutant variant carrying the truncating mutation p.Arg150stop (pCI-HA-R150X) were cloned into the pCI mammalian expression vector with an N-terminal HA tag (Figure S1). The wild-type cDNA fragment (1440 bp, encoding 470 amino acids) and the mutant fragment (477 bp, encoding the first 150 amino acids) were PCR-amplified from cDNA synthesized from HSkMCs, using specific primer pairs (Supplementary Table S1). A schematic overview of the cloning strategy is provided in Figure S1B and S1C. The ligated plasmids were transformed and positive colonies were selected and cultured for plasmid extraction using the QIAprep Spin Miniprep Kit (Qiagen). The integrity and orientation of the recombinant constructs were confirmed by Sanger sequencing using pCI vector-specific primers (Supplementary Table S1).

### Transient Overexpression of Wild-Type and Mutant Desmin Constructs in HSkMCs and AC16 Cells

Recombinant plasmids encoding wild-type desmin (pCI-HA-WT) and the R150X mutant variant (pCI-HA-R150X) were transiently transfected into primary human skeletal muscle cells (HSkMCs) and the AC16 cardiomyocyte cell line. Electroporation of HSkMCs was performed using the Neon Transfection System (Invitrogen) under the following conditions: voltage 1475 V, pulse width 20 ms, and 2 pulses. Transfection efficiency for both constructs in HSkMCs is shown in Figure S2. For AC16 cells, transient transfection was carried out using FuGENE HD Transfection Reagent (Promega corporation) following the manufacturer’s protocol.

### Lentivirus mediated Desmin R150X Mutant Cell Line Generation

For Cas9 and sgRNA expression in AC16 cells, we used the lentiCRISPR v2 vector in combination with lentiviral packaging plasmids to generate and deliver the CRISPR/Cas9 system (Supplementary Table S2). sgRNA sequences were designed using the CRISPOR design tool (Supplementary Table 1) and cloning plus lentiviral production were performed as per established protocols with minor modifications (24). sgRNAs were designed to target exon 1 of the DES gene in close proximity to the c.448C>T transition site to enable precise CRISPR-mediated modification (Figure S3). Briefly, plasmids were transfected into HEK293 cells and after 72 h of incubation, culture supernatants were collected, and viral particles were concentrated using Lenti-X™ Concentrator (Takara). Viral titers were estimated with Lenti-X™ GoStix® (Takara), and the required volume for the desired multiplicity of infection (MOI) was calculated. AC16 cells were transduced with lentiviral particles at an MOI of 4 in the presence of polybrene (8 ng/mL) to facilitate viral attachment. After 24 h, the medium was replaced with fresh medium containing puromycin (2 µg/mL) and incubated for 48 h for selection of transduced cells, which were subsequently replated for downstream experiments. These R150X mutant Desmin-expressing AC16 cells will hereafter be referred to as Des-R150X-CRISPR in the following sections.

### MG132 and Rapamycin Treatment

To assess the involvement of proteasomal degradation and autophagy in the clearance of the R150X desmin mutant, AC16 cells were treated with MG132 and Rapamycin. Both treatments were administered 6 hours prior to the end of each post-transfection time point (24 h, 48 h, 72 h, and 96 h). MG132 was used at a final concentration of 5 μM for 6 hours based on optimization studies (Figure S4A). Similarly, Rapamycin was applied at a final concentration of 50 nM for 6 hours in all subsequent experiments following standardization (Figure S4 B, C).

### Protein isolation and Western blotting

Protein lysates were prepared from both electroporated HSkMCs and transfected AC16 cells using lysis buffer containing 20 mM HEPES (pH 7.6), 20% glycerol, 100 mM NaCl, 1.5 mM MgCl₂, 0.2 mM EDTA, and 0.1% Triton X-100, supplemented with 1 mM DTT, 1 mM PMSF, and a protease inhibitor cocktail. Cell suspensions were sonicated on ice with three 10-second bursts. Total protein concentration was quantified using the Bradford assay to ensure equal sample loading. For western blotting, 50 μg of total protein was mixed with 2× Laemmli sample buffer boiled at 95 °C for 5 minutes, resolved on 12% SDS-PAGE, and transferred to nitrocellulose membranes. Membranes were blocked in 5% non-fat milk prepared in TBST buffer for 1 hour at room temperature. Blots were incubated with primary antibodies overnight at 4 °C, followed by HRP-conjugated secondary antibodies for 2 hours at room temperature. Protein detection was performed using an enhanced chemiluminescence (ECL) detection system (Thermo Fisher Scientific). β-Actin was used as loading control. The antibodies used are listed in Supplementary Table S3.

### Immunocytochemistry and Confocal Microscopy

Electroporated and transfected cells were fixed either with 4% paraformaldehyde for 15 minutes at room temperature or with cold acetone-methanol (1:1) for 5 minutes, depending on antibody-specific requirements. Following fixation, cells were permeabilized and blocked before primary antibody incubation. Coverslips were incubated with primary antibodies either overnight at 4 °C or for 1 hour at room temperature, as per the respective antibody guidelines, followed by a 1-hour incubation with secondary antibodies at room temperature. Nuclei were counterstained with DAPI for 5 minutes at room temperature. After washing, coverslips were mounted using ProLong™ Gold Antifade Mountant (Invitrogen), and imaging was performed using a Nikon Eclipse Ti2 confocal microscope. Confocal images were acquired using a 60× oil immersion objective, with excitation at 405, 488, 561, and 647 nm laser lines. For HSkMCs, glass coverslips were pre-coated with poly-L-lysine prior to electroporation. When mitochondrial staining was required, cells were incubated with 50 nM MitoTracker™ Red CMXRos in serum-free media for 30 minutes prior to fixation. The antibodies used are listed in Supplementary Table S4.

### Aggresomal detection Assay

Formation of aggregate-like structures were monitored by using Aggresomal Detection Kit (Abcam) that incorporates a 488 nm excitable, red-shifted molecular rotor dye that enables the specific detection of misfolded and aggregated protein species sequestered within aggresomal and aggresomal-like inclusions in fixed and permeabilized cells. The fluorescence associated with aggregated proteins was validated using both positive control (5uM, 6h MG132 treated cells) and negative control (Untreated as well as Untransfected cells) (Figure S5).

AC16 cells were transfected with both pCI-HA-WT and pCI-HA-R150X and further processing was done following manufacturer’s protocol. The coverslips were incubated with HA primary antibody for visualization of recombinant desmin and fluorescence signals corresponding to aggresomes were visualized using a standard Texas Red filter set.

### Image and Statistical Analysis

Regions of Interest (ROIs) were manually drawn for cells transfected with pCI-HA-WT and pCI-HA-R150, and Pearson’s correlation coefficient was calculated using Nikon AR software (version 5.0) across the entire z-stack of each transfected cell. The number of puncta was quantified using the ‘Analyze Particles’ function in Fiji (ImageJ) (25), and all densitometric analyses were also performed using Fiji. Statistical analysis was conducted using an unpaired t-test in GraphPad Prism (version 8.4.3), with p < 0.05 considered statistically significant.

## Results

### *R150* Desmin Mutation Leads to Premature Truncation and Perinuclear Aggregation in Myocytes

The stop gain mutation NM_001927.4: c.448C>T; p. (Arg150Stop) in the *DES* gene, introduces a premature stop codon within exon 1. This homozygous mutation was found in a patient born to consanguineous parents and presented with hallmark clinical features of desminopathy, including early-onset hypotonia, dilated cardiomyopathy (DCM), and granular deposits observed in skeletal muscle biopsy (22) —features consistent with myofibrillar myopathy (MFM) (Figure 1A–B). This C-to-T transition results in truncation of the desmin protein at the 150th amino acid (Arg150Stop), effectively eliminating the rod domain necessary for filament formation and cytoskeletal integration (Figure 1C). Multiple sequence alignment using Clustal Omega revealed that the Arg150 residue is highly conserved across species, underscoring its functional importance in desmin structure and filament assembly (Figure 1D).

**Figure 1:**
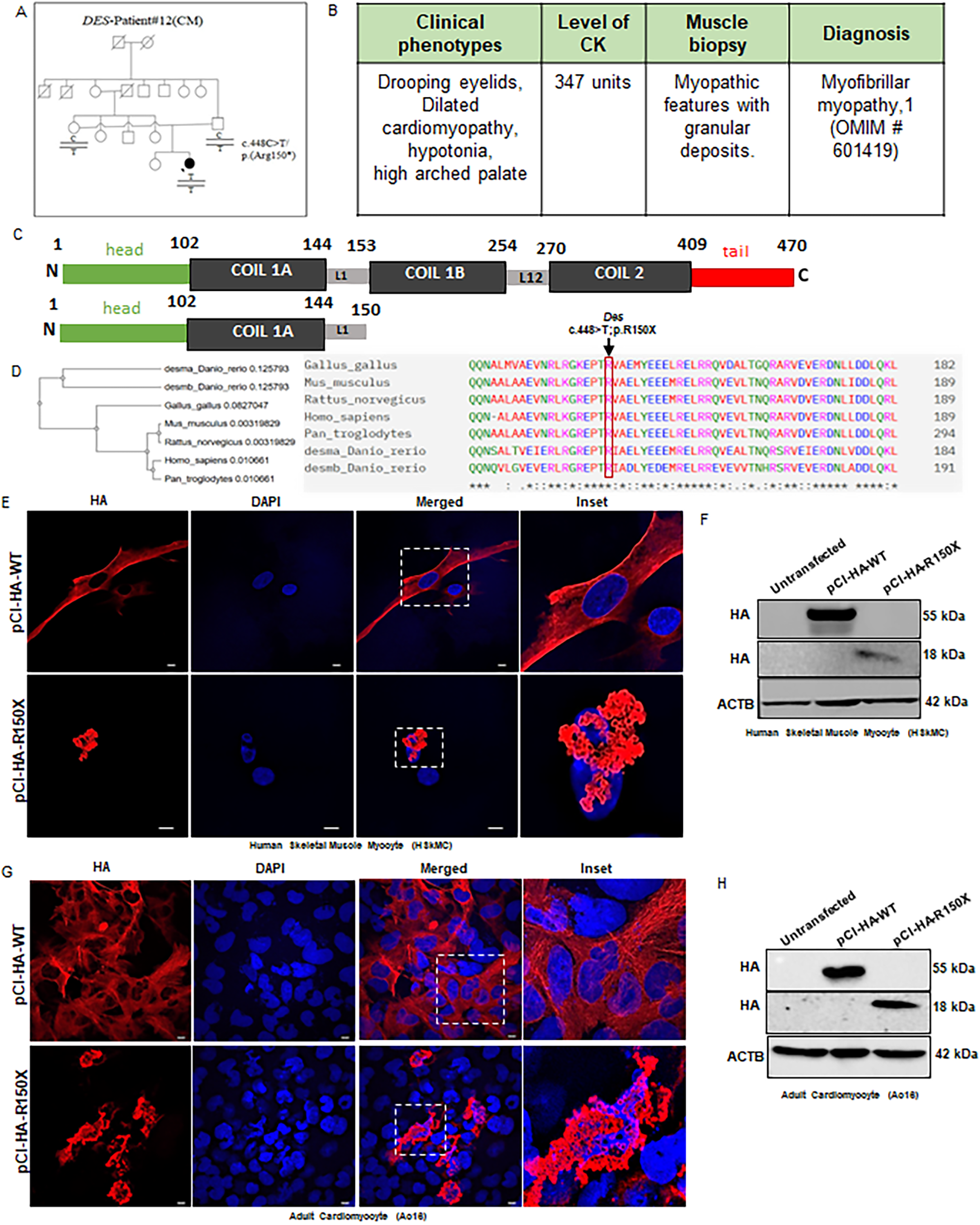
Characterization of the R150X Desmin mutation. (A) Segregation analysis confirming the presence of the R150X desmin mutation in the proband (16). (B) Clinical features of the proband carrying the R150X mutation. (C) Schematic representation of full-length desmin and the truncated protein formed due to R150X mutation. (D) Multiple sequence alignment showing that Arginine (R) at position 150 is highly conserved across species. (E) Confocal microscopy images (60X) of human skeletal muscle myocytes demonstrating perinuclear aggregate-like structures formed by R150X Desmin. (F) Western blot analysis confirming expression of the truncated 18 kDa protein in Human Skeletal Muscle Myocytes (HSkMCs). (G) Confocal microscopy images (60X) of Adult Cardiomyocytes showing perinuclear aggregate-like structures in R150X desmin–expressing cells. (H) Western blot analysis confirming the 18 kDa truncated protein in adult cardiomyocytes.

To investigate the functional consequences of this mutation, we generated constructs encoding HA-tagged wild-type (pCI-HA-WT) and mutant (pCI-HA-R150X) Desmin and overexpressed them in primary human skeletal myocytes (HsKMCs) and cardiomyocyte-derived AC16 cells. Immunofluorescence analysis demonstrated that cells expressing pCI-HA-WT formed an organized and filamentous desmin network extending from the nucleus to the cell periphery, indicating proper filament assembly (Figure 1E, G). In contrast, cells expressing pCI-HA-R150X exhibited a marked disruption in filament formation, instead showing intense accumulation of desmin-positive aggregate-like structures localized predominantly in the perinuclear region (Figure 1E, G). Western blot analysis confirmed the expression of an 18 kDa truncated desmin protein in R150X-expressing cells (Figure 1F, H). Our findings indicate that the R150X Desmin mutant results in a truncated protein that preferentially accumulates in the perinuclear region of both skeletal and cardiac muscle cells.

### Truncated R150 Desmin Disrupts Intermediate Filament Organization and Alters Cytoskeletal Architecture in Myocytes

Proper cytoskeletal organization is critical for myofibrillar architecture. Desmin forms a three-dimensional mesh around Z-discs, connecting them to the subsarcolemmal cytoskeleton, nuclei, and organelles to ensure mechanical integrity (1,26). As the truncated R150 mutant accumulates perinuclearly (Figure 1E, G), we hypothesized that this mutation may disrupt filament assembly and cytoskeletal organization. To investigate this, we transfected HsKMCs with either pCI-HA-WT or pCI-HA-R150X and performed co-immunostaining for different cytoskeletal elements including endogenous desmin (Figure 2A-F). In pCI-HA-WT transfected cells, endogenous Desmin exhibited the typical filamentous network similar to untransfected controls (Figure 2A). Moreover, both the endogenous desmin and the transfected wild-type desmin showed a complete colocalization (p<0.0001) in the transfected cells (Figure 2D). However, expression of R150X resulted in the collapse of this network, evident from loss of defined filamentous pattern, otherwise observed with endogenous desmin. (Figure 2A). Quantitative Pearson’s correlation analysis confirmed a significant reduction in colocalization between endogenous and mutant Desmin in R150X-expressing cells compared to wild-type (Figure 2D), indicating a dominant-negative effect of the mutant on filament integration.

**Figure 2:**
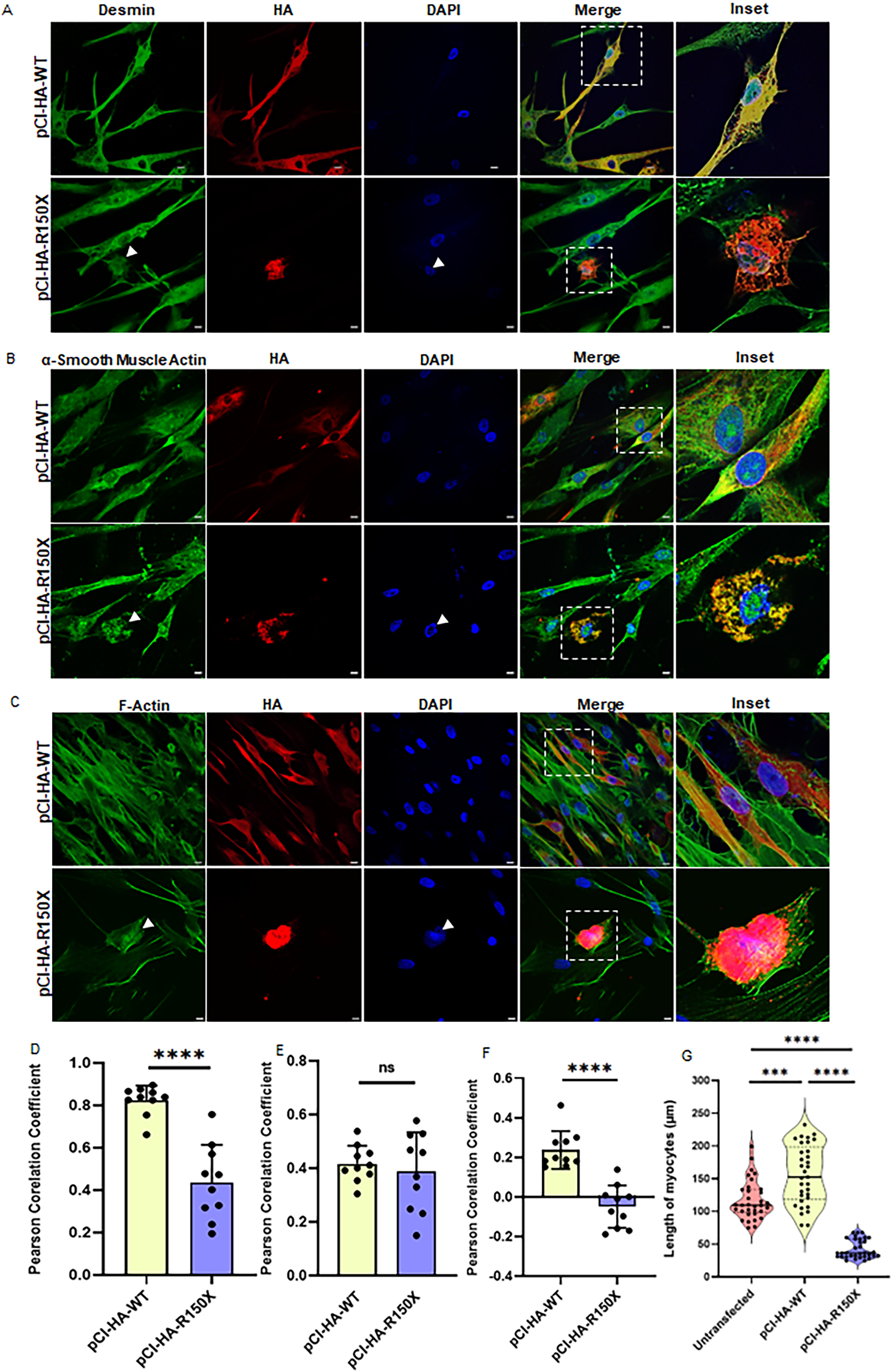
R150X Desmin expression disrupts endogenous desmin organization and alters cellular morphology. (A) Confocal microscopy images (60X) transfected with pCI-HA-WT and pCI-HA-R150X, showing HA-tagged Desmin (red) and endogenous Desmin (green). Expression of R150X leads to formation of aggregate-like structures and disruption of endogenous intermediate filament networks. (B) Distribution of α-smooth muscle actin (green) in pCI-HA-WT and pCI-HA-R150X transfected cells. Truncated desmin expression perturbs α-smooth muscle actin organization. (C) Distribution of F-actin (green) in pCI-HA-WT and pCI-HA-R150X transfected cells. R150X expression alters F-actin organization. (D) Pearson’s correlation coefficient analysis showing reduced colocalization between overexpressed HA-Desmin and endogenous desmin in R150X transfected cells (n=10, ****p<0.0001). (E) Pearson’s correlation coefficient analysis of HA-Desmin with α-smooth muscle actin show no change in colocalization between overexpressed HA-Desmin and α-smooth muscle actin in R150X transfected cells (n=10). (F) Pearson’s correlation coefficient analysis showing reduced colocalization between transiently expressed HA-Desmin and F-actin in R150X transfected cells (n=10, ****p<0.0001). (G) Quantification of myocyte length showing altered cellular morphology upon R150* expression. R150X transfected cells showed reduced myocyte length as compared to untransfected and wild-type expressing cells (n=33, ***0.0003, **** p<0.0001). All data are presented as the mean ± S.D.

We also observed alterations in gross cellular morphology, notably in cells forming aggregates (Figure 2A-C and G). To assess whether the R150X mutation impacts the broader cytoskeletal framework, we stained cells for α-smooth muscle actin (α-SMA) and filamentous actin (F-actin). Cells expressing pCI-HA-WT retained elongated morphology and showed well-organized, filamentous actin distribution consistent with normal cytoskeletal architecture (Figure 2B, C). In contrast, R150X-expressing cells displayed disrupted actin networks, with shortened filament length and reduced colocalization with desmin (Figure 2F, G). F-actin, which typically localizes at the cell periphery, showed poor spatial overlap with the R150X mutant. The mutant remained confined near the nucleus, further supporting impaired interaction between Desmin and actin filaments.

Additionally, we evaluated the distribution of actin-binding proteins, such as Plastin-1 (PLS1) and α-actinin-1 which are key regulators of actin filament cross-linking (Figure S6A, B). Although none of these actin-binding proteins colocalized with wild-type or mutant R150X desmin (Figure S6C, D), their altered distribution in R150X-expressing cells implied morphological disarray. Collectively, these findings suggest that the truncated R150X mutant forms aggregate-like structures that sequester endogenous desmin and perturb the structural alignment of actin filaments, ultimately compromising cytoskeletal integrity and myocyte morphology.

### R150 Desmin Mutation Disrupts Organelle Distribution and Spatial Organization in Myocytes

Desmin has reportedly been shown to maintain intracellular architecture by anchoring and spatially organizing key organelles such as mitochondria, the Golgi, and the sarcoplasmic reticulum (26). To investigate whether the truncated R150X disrupts this organization, we examined mitochondrial, Golgi, and ER markers in transfected myocytes. In cells transfected with pCI-HA-R150X, mitochondrial distribution was notably altered compared to both pCI-HA-WT and untransfected controls. Mitochondria are evenly dispersed along the filamentous axis of the cell. The cells overexpressing R150X showed distribution of mitochondria clustered abnormally near the perinuclear aggregates of the truncated desmin (Figure 3A). Further, immunostaining with Golgi markers—GM130 (cis-Golgi) (Figure 3B and D), GCC2 (trans-Golgi) (Figure S7), and p230 (trans-Golgi) (Figure S7)—revealed a significant reduction in Golgi signal intensity in R150X-expressing cells, suggesting impaired Golgi maintenance or positioning. Interestingly, co-localization analysis with Calnexin, the ER resident protein, indicated that the R150X aggregates partially overlap with ER structures (Figure 3C, E), raising the possibility of ER entrapment. Collectively, these findings suggest that the R150X Desmin mutant disrupts the intermediate filament network, thereby altering the spatial distribution and potentially the function of critical organelles in myocytes.

**Figure 3.**
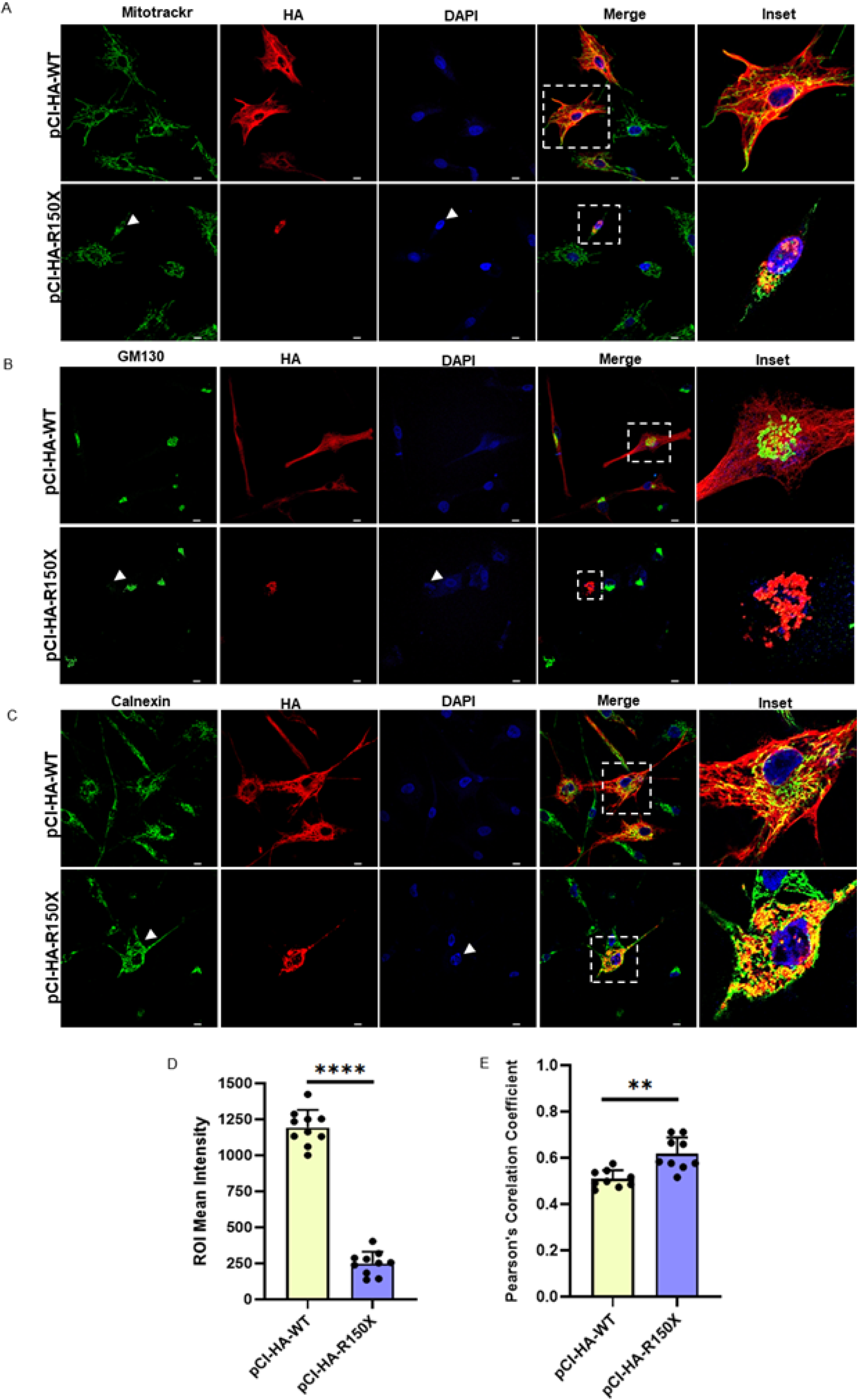
R150X desmin aggregate-like structures disrupt organelle distribution. (A) Confocal microscopy images (60X) transfected with pCI-HA-WT and pCI-HA-R150X, showing HA-Desmin (red) and MitoTracker (green). In R150X cells, mitochondria cluster around perinuclear aggregate-like structures. (B) Confocal microscopy images (60X) transfected with pCI-HA-WT and pCI-HA-R150X, showing HA-Desmin (red) and GM130, a cis-Golgi marker (green). In WT cells, Golgi localizes at the nuclear periphery, whereas R150X aggregate-like structures form large perinuclear clumps with loss of Golgi signal (arrows). (C) Confocal microscopy images (60X) transfected with pCI-HA-WT and pCI-HA-R150X, showing HA-Desmin (red) co-immunostained with Calnexin (green). R150X aggregate-like structures appear to colocalize with Calnexin. (D) Quantification of GM130 intensity showing a significant reduction in Golgi signal in R150X aggregate-harbouring cells (n=10, ****p<0.0001). (E) Pearson’s correlation coefficient analysis of HA-Desmin with Calnexin showing the truncated desmin colocalizing with Calnexin (n=9, **p=0.0052). All data are presented as the mean ± S.D.

### Truncated R150 Desmin Mutant Forms Persistent Aggregate-Like Structures in Cardiomyocytes

Several missense mutations in the *DES* gene have been previously reported to result in desmin misfolding and formation of intracellular aggregates or aggregate-like structures, which is a hallmark of desmin-related myopathies (11–15). To determine whether the truncated R150X mutant similarly forms aggregate-like structures, we transfected AC16 cardiomyocytes with pCI-HA-R150X and stained them using a red fluorescent molecular rotor dye specific for aggregated proteins. We observed prominent co-localizing foci between R150X and the aggresomal dye in transfected cells, which were absent in cells expressing pCI-HA-WT (Figure 4A). This finding was supported by a statistically significant increase in the Pearson’s correlation coefficient (p<0.0001) for colocalization between the mutant desmin and aggresome-specific dye (Figure 4B).

**Figure 4:**
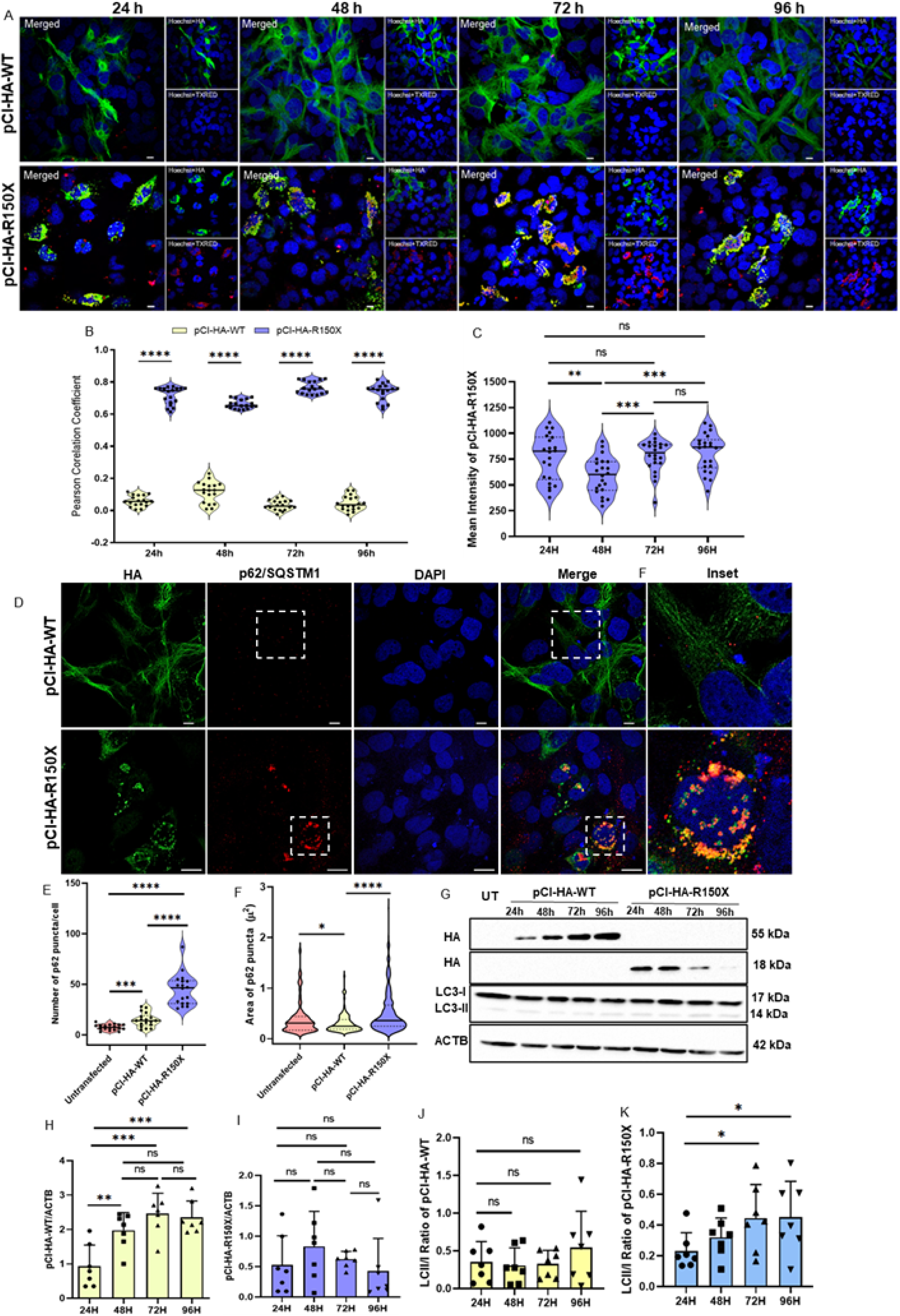
R150X aggregate-like structures show sustained accumulation with p62 recruitment and altered LC3 expression. (A) Confocal microscopy images (60X) of cells transfected with pCI-HA-WT and pCI-HA-R150X, showing aggregate-like structures in R150X-expressing cells detected by the aggresome detection assay. Overexpressed desmin is shown in the green channel, and the aggresome rotor dye in the Texas Red channel. (B) Quantification of Pearson’s correlation coefficient shows increased colocalization of overexpressed desmin (green) with the aggresome rotor dye (red) in cells expressing pCI-HA-R150X across all the four timepoints-24h,48h,72h and 96h (n=18, ****p<0.0001). (C) Quantification of mean intensity of overexpressed desmin (green) reveals that R150X aggregates persist without reduction over time (n=22, **p=0.0050[24v/s48], ***p=0.0005[48v/s72], ***0.0002[48v/s96]). (D) Confocal microscopy images (60X) of cells transfected with pCI-HA-WT and pCI-HA-R150X, shows that p62 colocalizes with R150X aggregates. (E) Quantification of puncta number per cell indicate increased p62 recruitment upon R150X expression (n=18, ***p=0.0003, ****p<0.0001). (F) Quantification of puncta area indicate increased enlargement of p62 puncta upon R150X expression (*p=0.0181, ****p<0.0001). (G) Western blot showing expression of wild-type and mutant R150X desmin under basal conditions. (H) Densitometric analysis of wild-type desmin at four different time-point depicts a gradual increase in the overexpression of pCI-HA-WT over time (n=7, **p=0.0050[24v/s48], ***p=0.0005[24v/s72], ***p=0.0004[24v/s96]). (I) Densitometric analysis of truncated desmin at four different time-point depicts no gradual change in the overexpression of pCI-HA-R150X over time (n=7). (J) Time-course analysis of LC3-II/I ratio in cells transfected with pCI-HA-WT under basal conditions depicts no such differences in LCII/I ratio over time (n=7). (J) Time-course analysis of LC3-II/I ratio in cells transfected with pCI-HA-R150X under basal conditions depicts an increase of LCII/I ratio over time (n=7,* p=0.0412 [24v/s72], * p=0.0452 [24v/s72]). All data are presented as the mean ± S.D.

To assess the temporal dynamics of aggregate formation and clearance, we examined transfected cells at 24-, 48-, 72-and 96-hours post-transfection. The R150X aggregate-like structures were persistent throughout the time course, suggesting that these misfolded structures are not efficiently cleared (Figure 4C). These observations indicate that the R150X mutant promotes stable aggregate-like structures in cardiomyocytes, highlighting a defect in the intrinsic protein quality control pathways responsible for maintaining desmin proteostasis.

### p62/SQSTM1 is Upregulated and Associates with R150X Aggregates-like structures in Cardiomyocytes

Aggregate-like structures often recruit p62/SQSTM1, a multifunctional ubiquitin-binding scaffold that links polyubiquitinated aggregates to autophagosomes for degradation (27,28). Elevated p62 levels have been reported in Desmin mutant models as a compensatory response to proteotoxic stress (29). To assess whether this mechanism is activated in response to the truncated desmin mutant, we assessed p62 expression in cardiomyocytes transfected with pCI-HA-R150X.

A statistically significant increase in the number and area of p62-positive puncta was observed in R150X expressing cells compared to cells transfected with full-length desmin (pCI-HA-WT) (Figure 4D-F). Notably, these p62-positive puncta were found in close spatial proximity to R150X aggregate-like structures, suggesting that p62 is actively recruited or sequestered with the misfolded protein structures. These observations indicate that p62/SQSTM1 accumulates around R150X aggregate-like structures as part of an adaptive proteostatic response. However, the continued presence of these aggregate-like structures despite p62 upregulation implies an insufficient or overwhelmed clearance system.

### Impaired Clearance of Truncated R150 Desmin Promotes Autophagic Induction

Misfolded or truncated proteins are typically eliminated from the cytoplasm via tightly regulated degradation pathways such as the ubiquitin-proteasome system (UPS), the autophagy-lysosome pathway routes to maintain cellular proteostasis and prevent toxic protein accumulation (30–32). To examine the clearance dynamics of the truncated R150X desmin mutant, we selectively inhibited the proteasomal degradation pathway using MG132, a potent and reversible 26S proteasome inhibitor that blocks UPS-mediated degradation and promotes accumulation of ubiquitinated proteins (33). We hypothesized that if the truncated R150X Desmin protein is being targeted for proteasomal degradation, its cellular accumulation should increase upon MG132 treatment. MG132 treatment led to a significant increase in R150 levels only at 24 h, while similar increases at 48, 72, and 96 h did not reach statistical significance (Figure S8).

Immunoblot analysis at different time points under basal conditions showed a decreasing trend in R150X expression over time, but these changes were not statistically significant, indicating that the mutant protein persists as aggregate-like structures (Figure 4A–C, G–I). Given this persistence, we next investigated whether autophagy was activated as a compensatory mechanism in response to proteasomal insufficiency by assessing LC3-II/I levels. A time-dependent increase in the LC3-II/I ratio was observed in pCI-HA-R150X– transfected cells, particularly at 72 h and 96 h, whereas no such trend was evident in pCI-HA-WT–transfected cells (Figure 4J, K). Notably, while proteasomal activity appeared sufficient to clear aggregates at 24 h, its capacity diminished over time, coinciding with activation of autophagy as a secondary clearance pathway. These findings suggest that persistent accumulation of R150X aggregates may induce autophagic pathways to compensate for impaired proteasomal degradation.

### Induction of Autophagy Facilitates the Clearance of R150 Desmin Mutant in Cardiomyocytes

Autophagy, a critical homeostatic mechanism, degrades long-lived proteins and clears aggregated or misfolded proteins that evade proteasomal clearance (29,34). Since the R150X desmin mutant formed persistent aggregates, resisted UPS-mediated degradation, and showed elevated p62/SQSTM1 levels, we tested whether autophagy induction could facilitate its clearance. To evaluate this, we treated AC16 cardiomyocytes expressing either pCI-HA-WT or pCI-HA-R150X with rapamycin, a well-characterized mTORC1 inhibitor and autophagy inducer, at four post-transfection time points: 24 h, 48 h, 72 h, and 96 h. Immunoblot analysis showed no change in pCI-HA-WT expression upon rapamycin treatment (Figure 5A–C). In contrast, truncated R150X levels consistently decreased across all time points following treatment, suggesting enhanced autophagy-mediated clearance (Figure 5D–F). However, LC3-II/I ratios remained unchanged in both wild-type and mutant conditions, regardless of rapamycin exposure (Figure 5C and F). Immunofluorescence analysis revealed a decrease in the intensity of pCI-HA-R150X and a decrease in both the number and area of p62-positive puncta following rapamycin treatment (Figure 5G-L). Although the intensity of aggregate-like structure formed by pCI-HA-R150X decreased significantly (Figure 5I), the signal intensity of the aggresome-specific rotor dye remained unchanged (Figure 5J). The decrease in both the number and area of p62-positive puncta following rapamycin treatment (Figure 5K and L) is consistent with increased autophagic flux and consumption of p62-bound aggregates. Collectively, these results suggest that activation of autophagy through rapamycin treatment facilitates the clearance of truncated R150X desmin mutant and may help restore cellular proteostasis in the context of desmin-related myopathies.

**Figure 5:**
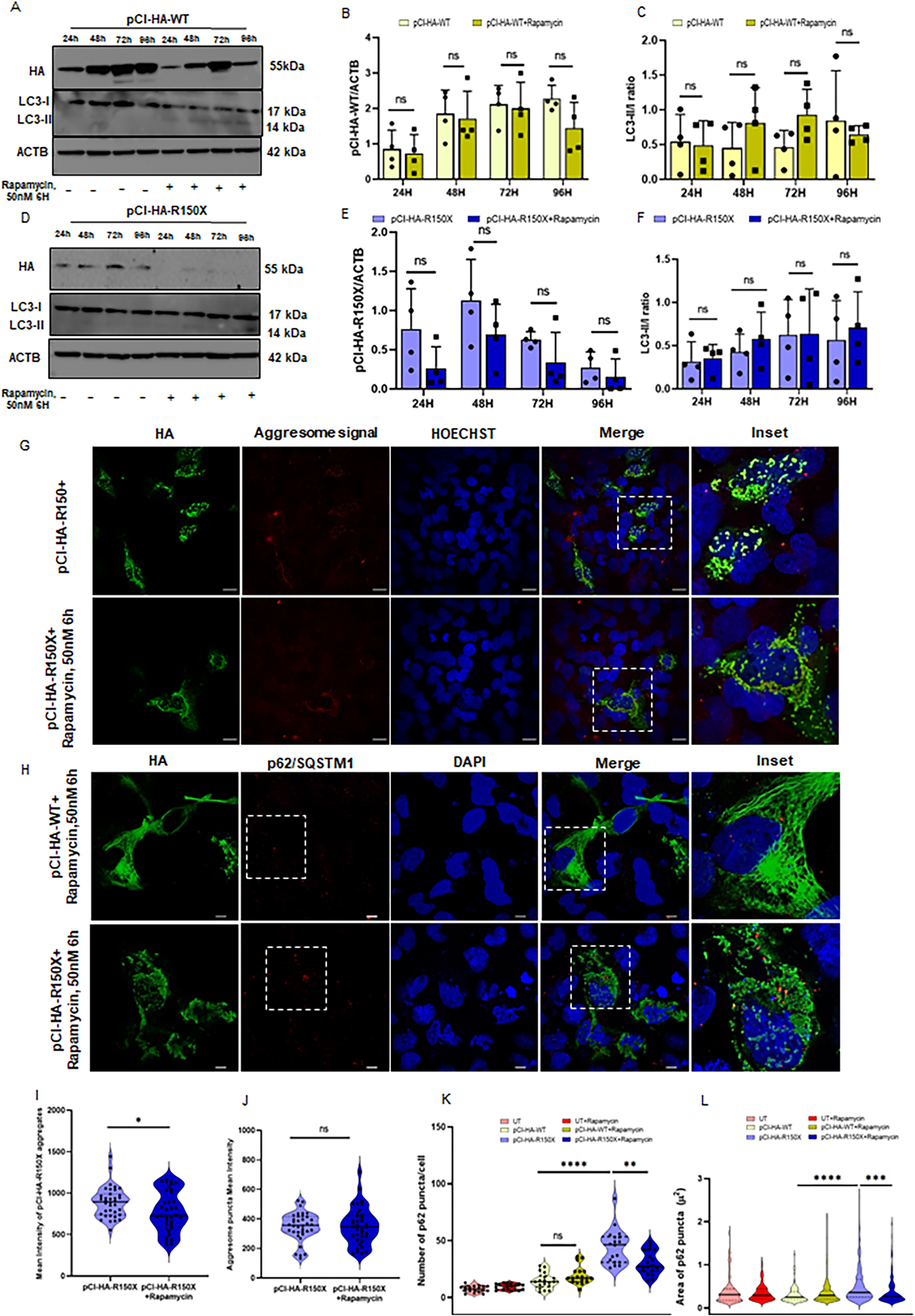
Rapamycin treatment reduces R150X desmin aggregate-like structures and modulates p62 expression. A) Western blot showing expression of wild-type desmin in the presence and absence of rapamycin (50nm, 6h). B) Densitometric analysis of pCI-HA-WT in presence and absence of rapamycin showed no change in expression of wild-type desmin levels upon rapamycin treatment (n=4). C) LC3-II/I ratio in cells expressing pCI-HA-WT in presence or absence of rapamycin treatment (n=4). D) Western blot showing expression of truncated R150X desmin in the presence and absence of rapamycin (50nm, 6h). (E) Densitometric analysis of R150X expression in presence and absence of rapamycin showed reduced expression of truncated R150X after rapamycin treatment (n=4). (F) LC3-II/I ratio in cells expressing pCI-HA-WT in presence or absence of rapamycin treatment (n=4). (G) Confocal microscopy images (60X) of cells transfected with pCI-HA-WT and pCI-HA-R150X, shows wild-type and mutant R150X desmin expression in presence or absence of rapamycin treatment. (H) Confocal microscopy images (60X) of cells transfected with pCI-HA-WT or pCI-HA-R150X, shows association of p62 with R150X aggregates after rapamycin treatment. (I) Quantification of mean intensity of overexpressed desmin (green) depicting reduction of R150X aggregates after rapamycin treatment (n=36, *p=0.0327). (J) Quantification of mean intensity of aggresome rotor dye intensity (red) shows no difference in intensity after rapamycin treatment (n=36). (K) Quantification of p62 puncta number per cell, showing decreased p62 accumulation in R150X transfected cells after rapamycin treatment (n=18, **p=0.0025). (L) Quantification of p62 puncta area, showing reduction in size after rapamycin treatment (***p=0.0006). All data are presented as the mean ± S.D.

### Endogenous R150X Cardiomyocyte Model Mirrors Overexpression-Induced Aggregation

Complete loss of desmin abrogates the formation of desmin-positive inclusions, as knockout models fail to develop the aggregates characteristic of desmin-related myopathies (19,20), supporting that inclusions arise from mutant or misfolded desmin rather than absence of the protein. In this context, desmin deficiency may bypass aggregate nucleation compared to aggregate-forming conditions which typically alter protein quality-control dynamics and autophagic response. To experimentally address this, we engineered the R150X pathogenic variant in AC16 cardiomyocytes using CRISPR/Cas9, targeting exon 1 near the c.448C>T transition to mimic the phenotype observed in our overexpression models (Figure 6A).

**Figure 6:**
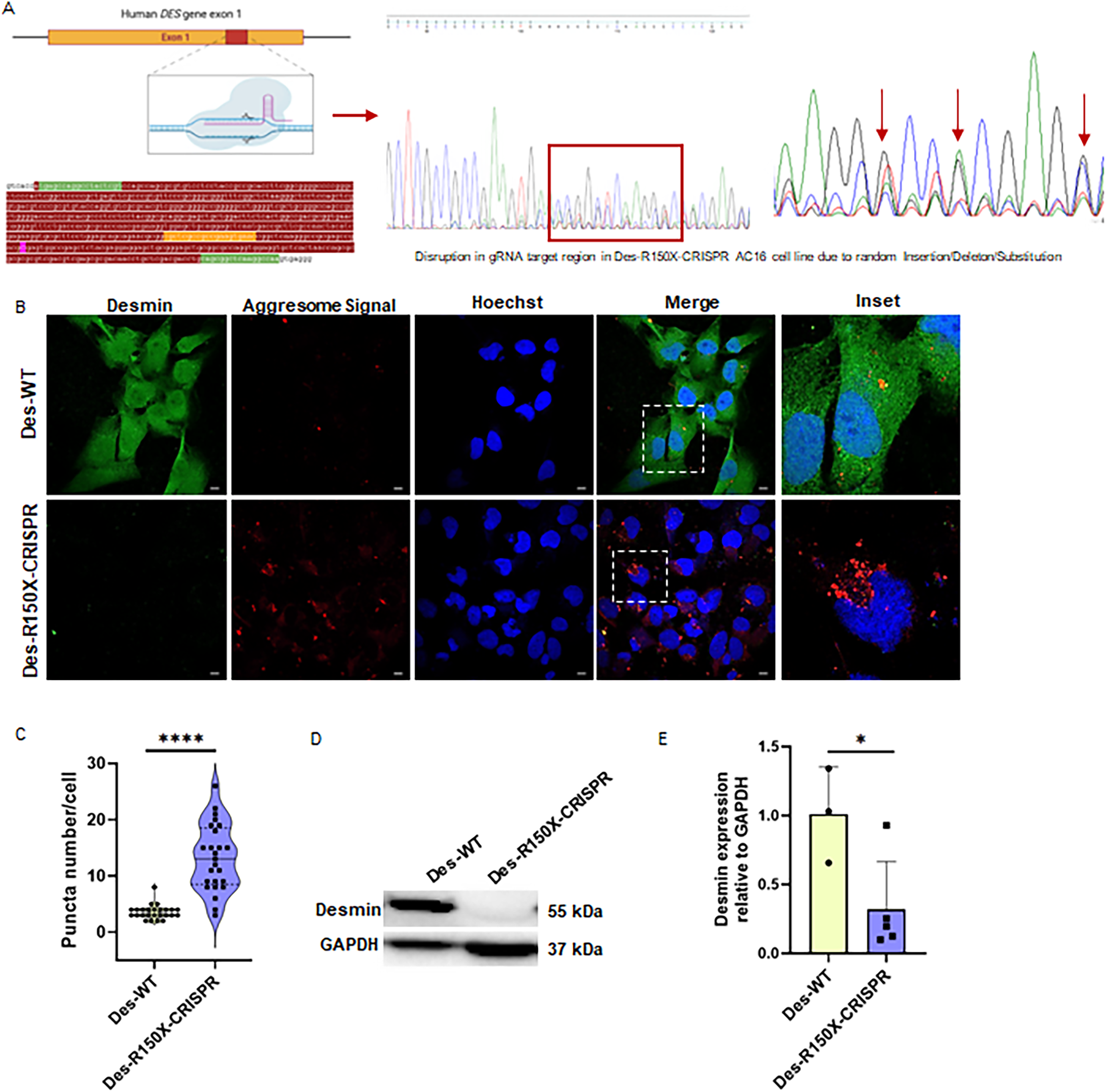
Characterization of Des-R150X-CRISPR cardiomyocyte model. A) Schematic representation of the target region in exon 1 of the DES gene showing the gRNA binding site. Sanger sequencing traces confirm the successful introduction of insertions, deletions, and substitutions at the target locus. B) Confocal microscopy images (60X) depicting increased aggresome rotor dye (red) signal in the Des-R150X-CRISPR cell line. C) Quantification of aggresome puncta per cell depicting formation of aggregate-like structures in the Des-R150X-CRISPR cell line (n=25,****p<0.0001). D) Western blot showing reduced desmin expression in the Des-R150X-CRISPR cell line as compared to Des-WT. E) Densitometric analysis confirming decreased desmin levels in the Des-R150X-CRISPR cell line (n=5, *p=0.0338). All data are presented as the mean ± S.D.

**Figure 7.**
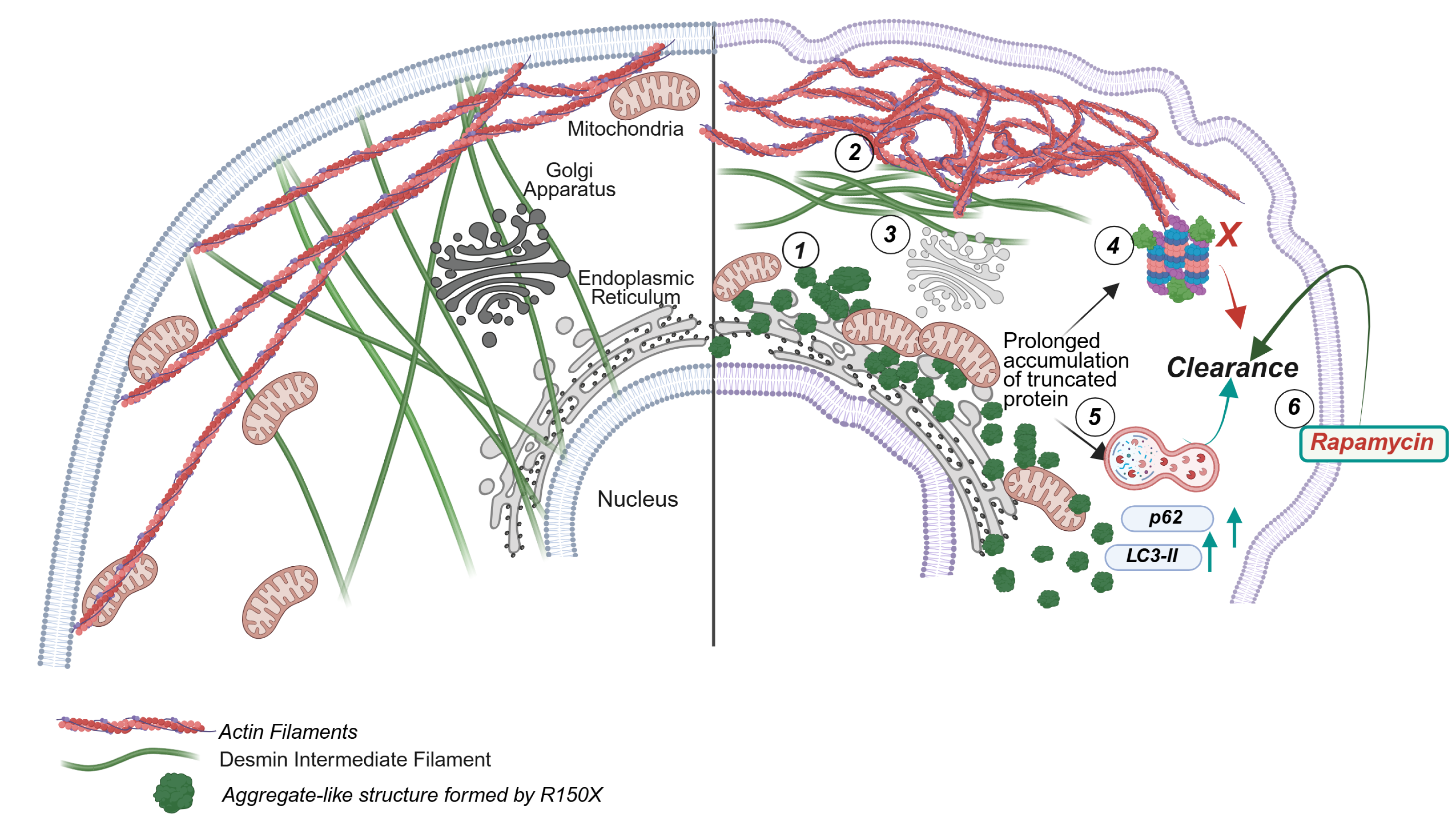
Schematic summary of the pathogenic effects of the R150X desmin mutant on structural integrity and cellular homeostasis. (1) R150X forms perinuclear aggregate-like structures. (2) Endogenous desmin, actin, and actin-binding protein distribution is disrupted. (3) Aggregate-like structures seems to be colocalizing with the endoplasmic reticulum and perturb Golgi and mitochondrial distribution. (4) Persistent accumulation of aggregate-like estructures overwhelms the proteasomal degradation pathway. (5) Key autophagic markers are upregulated as a compensatory response. (6) Pharmacological induction of autophagy via rapamycin facilitates aggregate clearance in myocytes

Quantitative confocal imaging revealed a significant increase in aggresome signal, as evident from elevated puncta numbers in Des-R150X-CRISPR compared with Des-WT (Figure 6B and C). Western blot analysis also confirmed a significant reduction in endogenous desmin levels (Figure 6D, E). The detection of the truncated endogenous desmin protein was not possible with the antibody used, as it targets the C-terminal epitope absent in the truncated form. These findings demonstrate that the formation of persistent aggregate-like structures and their downstream effects on cardiomyocyte architecture and homeostasis are not restricted to overexpression systems. Through precise genome editing mimicking the pathogenic c.448C>T variant in an endogenous context, we successfully recapitulated the aggregate-associated phenotype observed in the overexpression model likely responsible for the granular deposits reported in the muscle biopsy of the patient (16).

## Discussion

### Novel pathogenic DES R150X mutation as a model for Desminopathies

Desminopathies, particularly myofibrillar myopathies, are characterized by dense desmin-positive aggregates in skeletal and cardiac muscle, a feature consistently observed in cell and animal models (11–15,29). While many pathogenic DES mutations have elucidated how mutant desmin disrupts cellular architecture and homeostasis, most studies focus on single-point mutations or deletions, leaving truncating mutations largely unexplored. The novel DES R150X mutation, identified in a 13-year-old girl from consanguineous parents, presents a distinct desminopathy phenotype with dilated cardiomyopathy and distal upper-limb weakness (22). Muscle biopsy from the same patient published in another study revealed fiber-size variation, irregular Gömöri trichrome staining, increase in NADH-TR activity, and ultrastructural abnormalities including clustered lobulated nuclei and disrupted myofilaments with expanded Z-band aggregates, corroborating with histopathological features that are typical of desminopathies (23). Despite these findings, the study did not elucidate the mechanistic basis of this truncating mutation in disease pathogenesis. Complementary immunohistochemistry and western blot analysis also demonstrated a complete absence of desmin expression. In contrast, our findings highlight that the C-to-T transition at nucleotide 448 produces a truncated desmin protein that retains the capacity to form aggregate-like structures, suggesting a toxic gain-of-function rather than simple loss of expression. Using both overexpression systems and a CRISPR/Cas9-engineered R150X cardiomyocyte model, we demonstrate that this truncated protein is structurally compromised, aggregation-prone, and capable of disrupting the intermediate filament network—likely driving the observed myopathic phenotype. Given its reproducible aggregation behaviour, disruption of filament organization, and recapitulation of hallmark histopathological features *in vitro*, the R150X mutation represents a powerful and physiologically relevant model for elucidating desminopathy mechanisms.

### R150X disrupts intrinsic intermediate filament assembly and disorganizes myoskeletal organization and organellar positioning

Intermediate filaments (IFs) are essential for skeletal muscle integrity, interconnecting Z-discs, myofibrils, and linking them to the sarcolemma, nucleus, and mitochondria to maintain mechanical strength and sarcomere organization (35,36). The DES R150X truncation at amino acid 150 abolishes the C-terminal domain, critical for filament bundling, higher-order organization, and docking of IF-associated proteins (37–39). Loss of interactions with chaperones such as CRYAB and structural regulators like MTM1 likely promotes aberrant filament self-association and aggregate formation (35,40,41). Moreover, desmin aggregates can impair actin filament organization (15,42), as observed with actin-binding partners including Nebulin, Nebulette, and Z-disc–associated proteins plectin 1d, syncoilin, and synemin, whose interactions with desmin’s C-terminal are essential for actin network integrity (43–46). Our immunocytochemistry data corroborate this, showing that truncated desmin disrupts actin filament organization around the nucleus, reflecting progressive loss of cytoskeletal coordination.

Disruption of desmin IFs impairs mitochondrial distribution and dynamics, compromising energy homeostasis and contractile function (47,48). Patient data also indicated mitochondrial displacement without overt morphological defects and a selective ∼20% deficiency in complex IV activity (23), which aligns with observations in our R150X overexpression model. Additionally, we observed reduced Golgi fluorescence at the nuclear periphery in R150X-expressing cells. The Golgi, typically positioned via microtubule and actin attachments (49–52), may be affected due to disrupted actin architecture due to desmin aggregation. However, the precise mechanism underlying the reduced Golgi fluorescence intensity observed in R150-expressing cells remains to be elucidated. Additionally, it is also unknown if the accumulation at ER-exit sites impairs ER-to-Golgi transport, leading to retention of secretory cargos and delayed Golgi processing. Truncated desmin aggregates partially associate with the ER protein Calnexin, suggesting potential ER involvement. However, their impact on ER structure and dynamics remains unclear, as reported for another intermediate filament, vimentin (53). The R150X aggregates might also affect calcium flux, as supported by studies in Desma knockout zebrafish, which show unaltered muscular morphology but altered calcium flux (54). Overall, while R150X aggregates appear to influence organelle distribution and function, the underlying mechanisms require further investigation.

### Persistent aggregate-like structure formed by R150X induces selective autophagy to clear off the truncated protein

We found that truncated R150X formed persistent aggregate-like structures of varying number, morphology, and distribution, often clustering perinuclearly. Aggresome detection assays confirmed their aggresomal association, though the full complement of associated proteins remains unknown. Misfolded proteins, such as this truncated variant, are inherently less stable due to improper folding and the exposure of hydrophobic residues, which predispose them to degradation via protein quality-control pathways (55). While individual mutant molecules are unstable, their propensity to self-associate can drive the formation of persistent, aggregate-like structures that are resistant to clearance (56). One such candidate which we focused on this study is p62/SQSTM1, which facilitates aggresome formation and activates autophagy to protect cardiomyocytes from proteotoxic stress (28,29). Through its UBA domain, p62 recruits misfolded proteins; the PB1 domain drives oligomerization, and the LC3-interacting region (LIR) directs autophagic degradation, effectively linking the UPS and autophagy pathways (28).

Earlier studies suggest inefficient UPS clearance, could be potentially due to impairment of the 19S proteasome subunit in desminopathic mouse hearts (10). As the 19S subunit channels substrates into the 20S core, dysfunction could contribute to desminopathy pathogenesis. Under stress, excess abnormal proteins can overwhelm the UPS, impair proteasome activity, and activate the autophagy–lysosome pathway to relieve proteotoxic stress (57). In R150X mutants, LC3-II/I ratio increased but remained insufficient to clear aggregates. Similar autophagy defects occur in LMNA^–/–^ mice, where mTORC1 hyperactivation impairs autophagy; rapamycin reduces mTORC1 signaling, improves muscle function, and extends lifespan (58). Rapamycin treatment reduced R150X aggregate-like structures, yet LC3-II/I ratio remained unchanged. Although rapamycin activates autophagy via mTORC1 inhibition, LC3-II responses are context-dependent and may not always increase (59,60). Together, these findings suggest that while our results are consistent with enhanced autophagy, a more detailed analysis of flux would help strengthen this interpretation.

Working with a transient overexpression system presents limitations, including variability in pCI-HA-R150X expression upon rapamycin treatment. While the decreasing trend of R150X expression were consistently observed, the magnitude varied across replicates, resulting in minimal statistical significance (Figure 5B). This essentially reflects biological heterogeneity inherent to overexpression systems rather than an absence of the effect. This also highlights the need for robust cell and animal models to dissect how truncating desmin variants disrupt degradation pathways and drive disease pathogenesis. While C2C12-based models expressing mutant desmin (61) and ES cell systems with head-domain deletions (62) have been developed, CRISPR-engineered muscle cell model harbouring truncating desmin mutation has never been reported. Our study addresses this gap by demonstrating aggregate-like structure formation and its functional implications in a Des-R150X CRISPR cardiomyocyte line, corroborated by findings from our overexpression model. A key limitation of this model is that, due to desmin’s essential role in muscle cells, viable homozygous R150X CRISPR clones could not be generated, as multiple attempts led to loss of cell viability. Consequently, all experiments were conducted in a heterozygous background. This aligns with previous studies showing that complete loss of desmin causes lethality in mouse and human models (19,20). Similarly, heterozygous patients and mice initially retain near-normal muscle structure, but cumulative proteotoxic stress from mutant desmin leads to late-onset pathology (6,63) (Goldfarb & Dalakas, 2009; Clemen et al., 2013). This also highlights that the pathogenic mechanism of desmin-related myopathies, in which severe manifestations are frequently associated with autosomal recessive inheritance and are progressive, ultimately leading to fatal outcomes in affected patients. We also admit that our study is limited by its in vitro focus and does not yet fully resolve how organelle dysfunction and aggregate persistence translate into tissue-level pathology in vivo. Nevertheless, this system provides a physiologically relevant platform to deepen mechanistic insights into desminopathy pathogenesis and lays the groundwork for therapeutic exploration.

## Acknowledgements

This work was supported by Intramural funding of BRIC-National Institute of Biomedical Genomics to Moulinath Acharya. Sukanya Mitra is a recipient of a Senior Research Fellowship (No. 09/1033(12191)/2021-EMR-I) from the Council of Scientific and Industrial Research (CSIR), Ministry of Science and Technology, Govt. of India. The authors appreciate initial help and discussion with Dr. Kaushik Sengupta, Saha Institute of Nuclear Physics, Kolkata. The authors also thank Dr. Arnab Gupta, Indian Institute of Science and Education Research, Kolkata for providing us with GCC2 antibody.

## Author Contributions

SM designed and performed the experiments, analyzed the data including statistical interpretation, and drafted the manuscript. TG and AKM generated the lentivirus-mediated CRISPR-Cas9 R150X AC16 cell line. SS generated recombinant plasmids and provided preliminary data. MM supervised the experiments, guided data interpretation, and reviewed the manuscript. MA conceived the study, supervised the project, and drafted the manuscript.

