## Supplementary Material for "Novel truncating Desmin mutation Arg150Stop disrupts structural integrity and cellular homeostasis by formation of persistent aggregate-like structure"

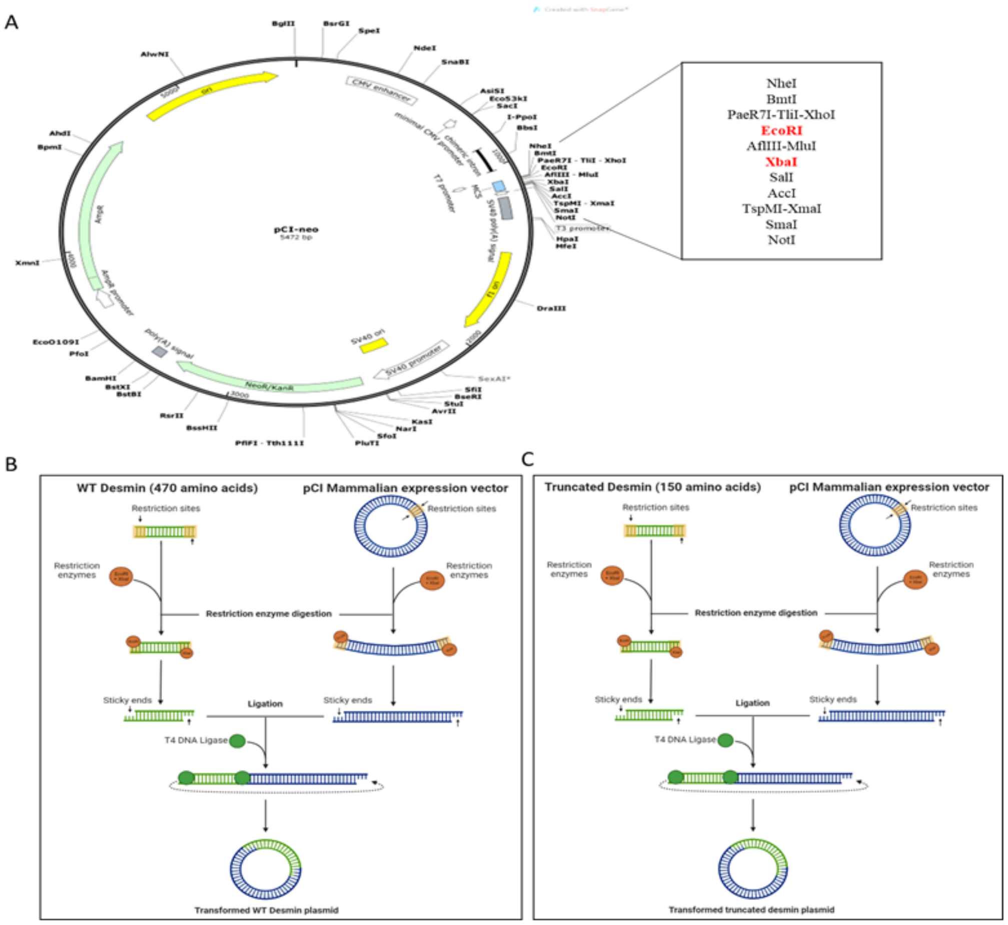


Figure S1: A) pCI Mammalian Expression Vector with restriction enzyme used for cloning full length wild-type (pCI-HA-WT) and mutant (p.Arg150stop; R150X) desmin variant (pCI-HA-R150X). B, C) The schematic representation of cloning of Wild-type (full-length) and R150X mutant (truncated) Desmin respectively

| **Primer ID** | **Primer sequence (5’ – 3’)** | **Regions Amplified (Product size in bp)** | **$T_A$ (ºC)** |
|---|---|---|---|
| DES-CL-F1 | CGGAATTC ATGTACCCATACGATGTTCCAGATTACGCTAGCCAGGCCTACTCGTCC | WT Desmin cDNA sequence (1440) | 64 |
| DES-CL-R1 | GCTCTAGATTAGAGCACTTCATGCTGCTG | | |
| DES-CL-F1 | CGGAATTC ATGTACCCATACGATGTTCCAGATTACGCTAGCCAGGCCTACTCGTCC | Truncated Desmin cDNA sequence (477) | 64 |
| DES-CL-R2 | GCTCTAGATCACGTCGGCTCGCGGCCCTT | | |
| pCI-T7-FP | TAATACGACTCACTATAGGG | pCI specific forward primer T7-promoter for sanger sequencing | |
| EBV-RP | GTGGTTTGTCCAAACTCATC | pCI specific reverse primer for sanger sequencing | |
| Des_sgRNA Forward | CACCGcgctcgccgccgaagtgaac | Lentivirus mediated Desmin R150X Mutant Cell Line Generation | |
| Des_sgRNA Reverse | AAACgttcacttcggcggcgagcgC | Lentivirus mediated Desmin R150X Mutant Cell Line Generation | |

Table S1: List of primers used

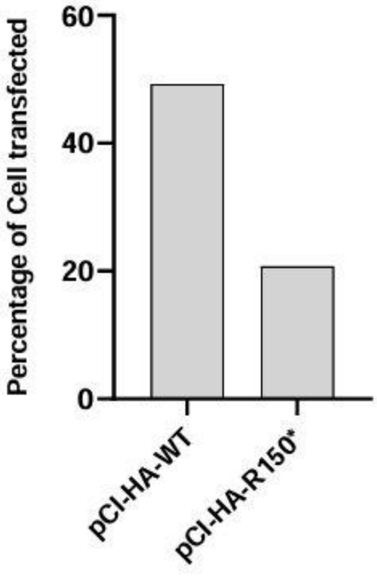


Figure S2: Transfection Efficiency of pCI-HA-WT and pCI-HA-R150X using Neon Transfection System (Invitrogen).

| Plasmid | |
|---|---|
| psPAX2 ( Addgene Plasmid #12260) | 2nd generation lentiviral packaging plasmid |
| pMD2.G( Addgene Plasmid #12259) | VSV-G envelope expressing plasmid |
| lentiCRISPR v2<br>(Addgene Plasmid #52961) | Lentiviral transfer plasmid, also known as pLentiCRISPR v2, for CRISPR/Cas9 genome editing of mammalian cells with a single guide RNA (sgRNA) |

Table S2: List of plasmids used for Lentivirus mediated Desmin R150X Mutant Cell Line Generation

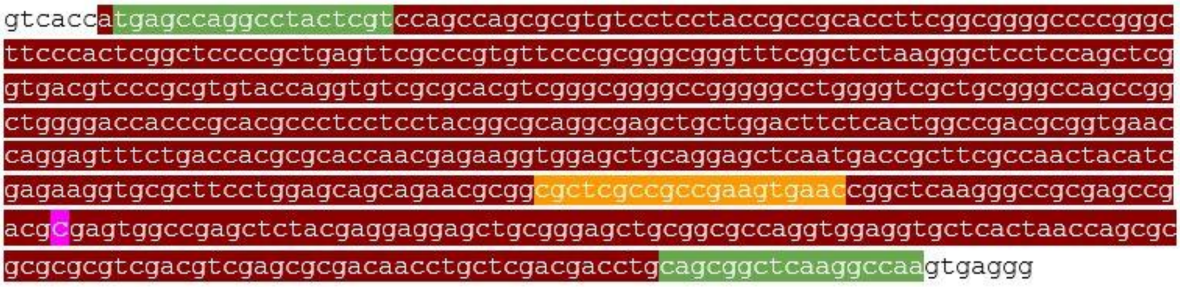


Figure S3: Sequence of exon 1 of the DES gene showing the c.448C nucleotide (highlighted in purple) and thecorresponding sgRNA binding site (highlighted in orange).

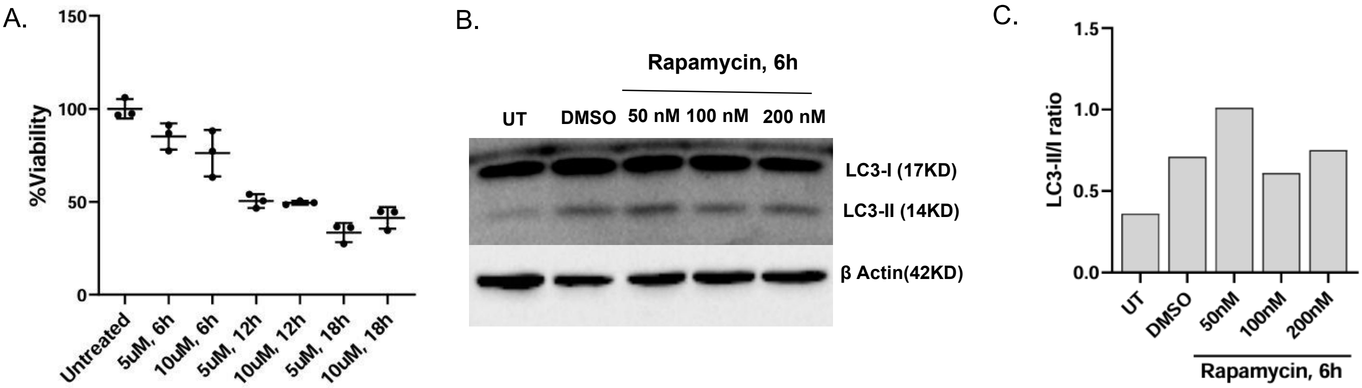


Figure S4: (A)The optimal dose and treatment duration of MG132 were standardized based on cell viability measurements using the MTT assay. (B, C) Standardization of the optimal concentration of Rapamycin by accessing LCII/LC3-I ratio via immunoblotting.

| Antibody used in Western Blot | Identifier | Dilution |
| --- | --- | --- |
| HA-Probe (F-7), Mouse monoclonal IgG2a | Santa Cruz Biotechnology | 1:1000 |
| Rabbit Anti-Desmin antibody, Source no- 21190 | Sigma-Aldrich | 1:1500 |
| polyclonal Rabbit LC3B | MBL International | 1:1000 |
| ACTB; Mouse monoclonal (C4) | Santa Cruz Biotechnology | 1:1000 |
| Goat Anti-Rabbit IgG H&L | Abcam | 1:10,000 |
| Goat Anti-Mouse IgG H&L | Abcam | 1:10,000 |

Table S3: List of antibodies used for western blot

| Antibody used in Immunofluorescence | Identifier | Dilution |
| --- | --- | --- |
| HA-Probe (F-7), Mouse monoclonal IgG2a | Santa Cruz Biotechnology | 1:200 |
| HA-Tag (C29F4) Rabbit mAb | Cell Signalling Technology | 1:400 |
| Rabbit Anti-Desmin antibody, Source no- 21190 | Sigma-Aldrich | 1:20 |
| mouse monoclonal anti-alpha smooth muscle actin antibody | Abcam | 1:200 |
| rabbit polyclonal anti-PLS1 antibody | Abcam | 1:200 |
| rabbit polyclonal to anti-alpha actinin/ACTN1 antibody | Abcam | 1:50 |
| Purified Mouse anti-GM130 | BD Transduction Laboratories™ | 1:200 |
| Rabbit polyclonal GCC2 | Altas Antibodies] | 1:500 |
| Purified Mouse Anti-Human p230 trans Golgi | BD Transduction Laboratories™ | 1:1000 |
| mouse Monoclonal Calnexin (AF18) | Santa Cruz Biotechnology | 1:200 |
| mouse monoclonal anti-SQSTM1 / p62 antibody [2C11] | Abcam | 1:200 |
| MitoTracker™ Red CMXRos | Cell Signalling Technology | 50nM |

Table S4: List of antibodies used for Immunofluroscence

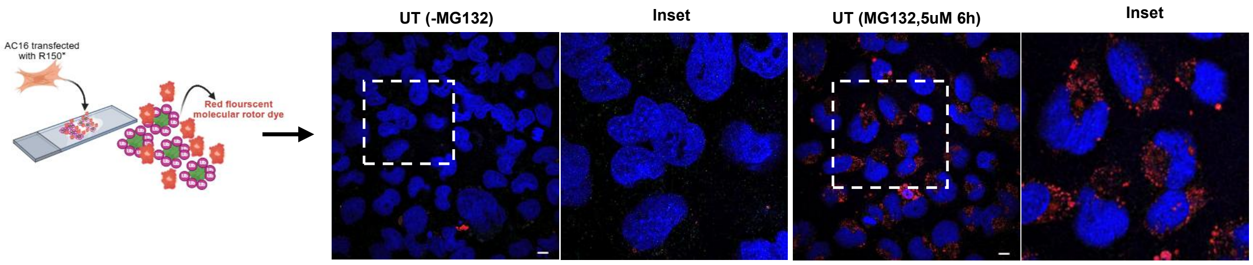


Figure S5: Validation of aggresomal fluorescence detection using control treatments. UT (-MG132) is negative control and UT (MG132, 5uM, 6h) is positive control.

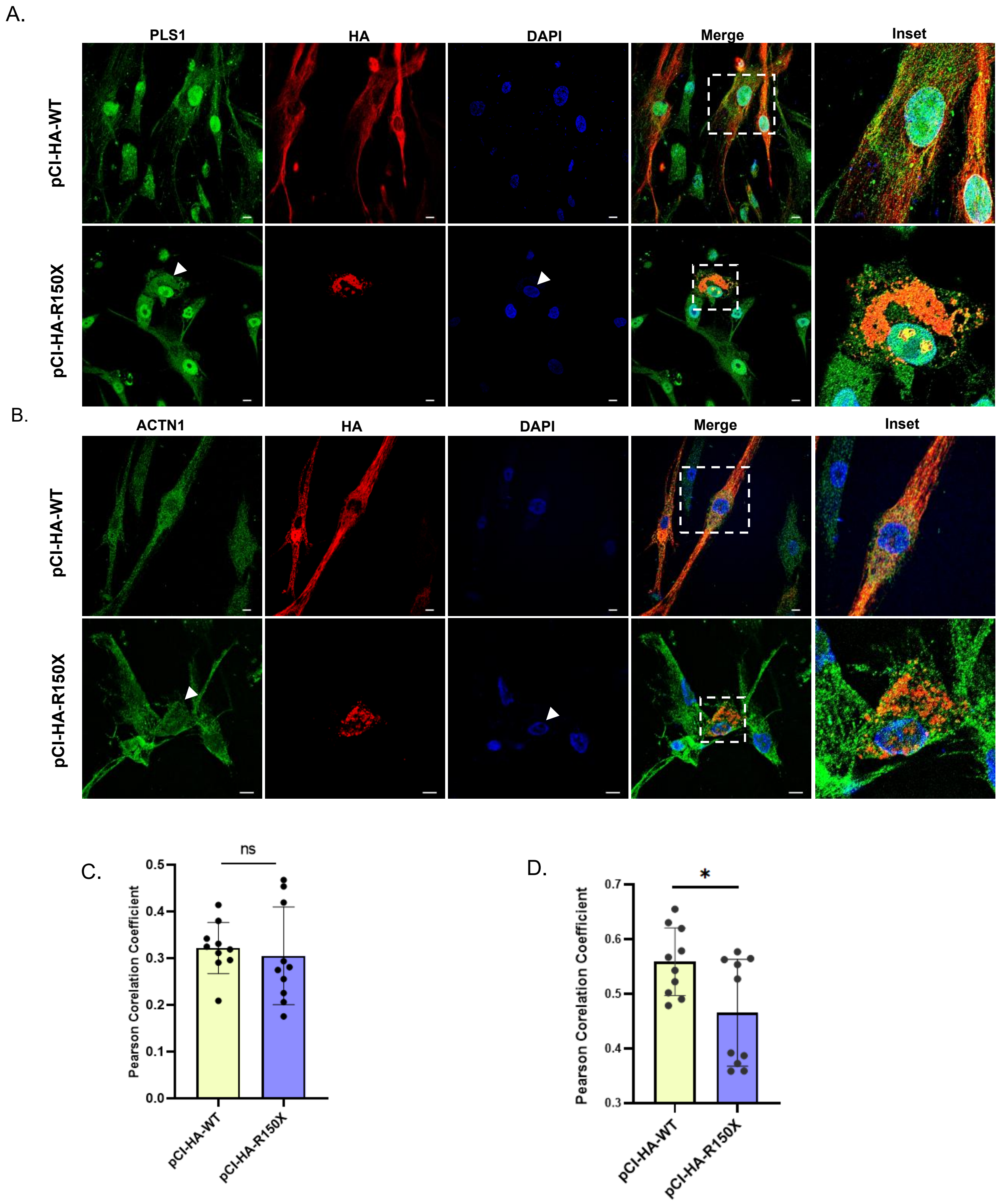


Figure S6: The distribution of Actin-binding proteins was perturbed in cells expressing truncated desmin depicted that pCI-HA-R150X affected the morphology of skeletal muscle cells.

A,B. pCI-HA-WT and pCI-HA-R150X transfected cells expressing HA tagged desmin (red) and co-stained for endogenous PLS1 and ACTN1 respectively (green)

C,D. Pearson corelation coefficient of transiently expressed desmin and endogenous PLS1 and ACTN1 respectively.

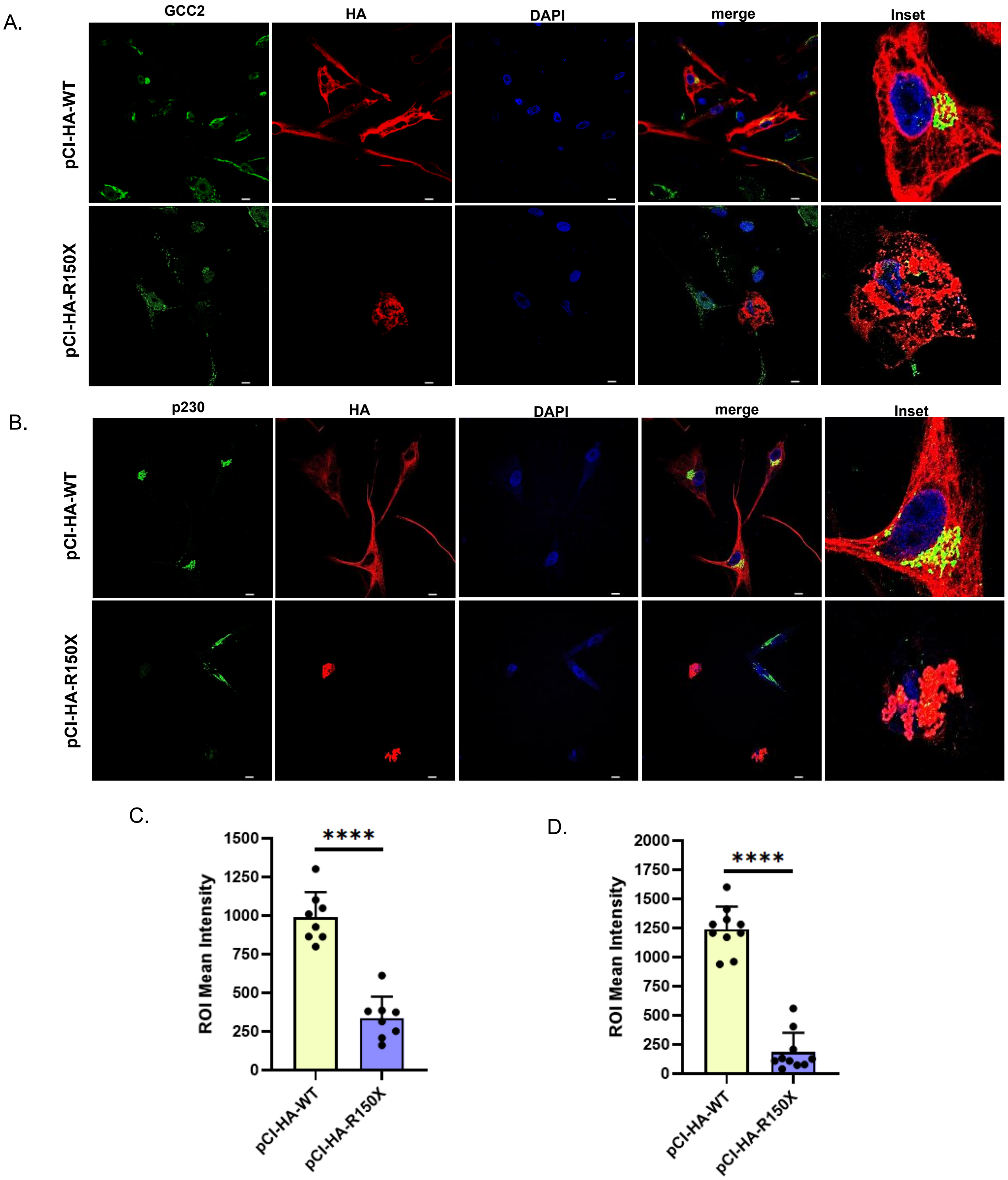


Figure S7: The expression of trans-golgi markers were also perturbed in cells expressing pCI-HA-R150X

A,B. pCI-HA-WT and pCI-HA-R150X transfected cells expressing HA tagged desmin (red) and co-stained for endogenous GCC2 and p230 respectively (green)

C,D. ROI mean Intensity of transiently expressed desmin and endogenous GCC2 and p230 respectively

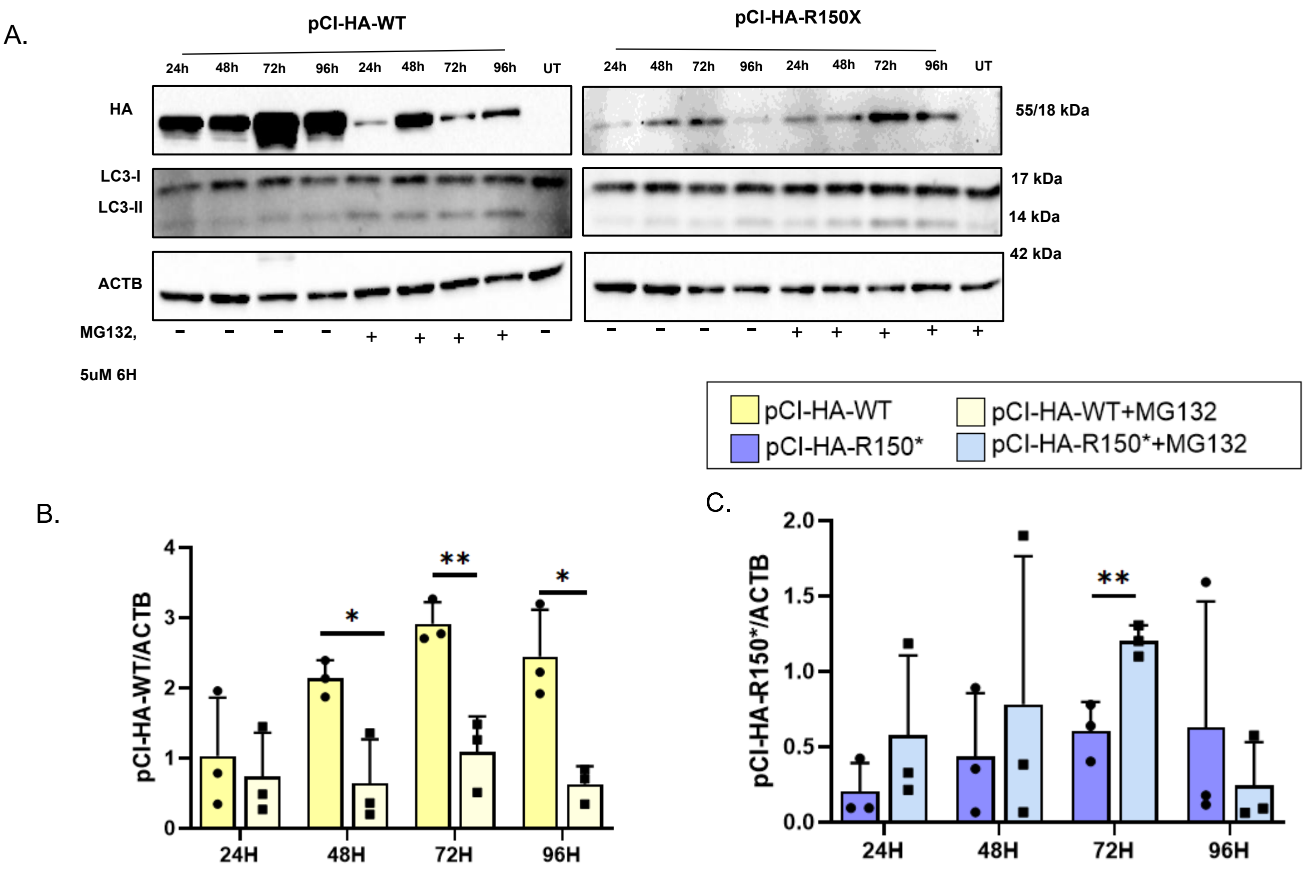


Figure S8: Western blot showing expression of wild-type and truncated R150X desmin in the presence and absence of MG132 (5µm, 6h).

A. Western blot depicting the expression level of wild-type and mutant R150X in presence and absence of proteasomal inhibitor MG132
B. Quantification of wildtype expression in presence and absence of proteasomal inhibitor MG132
C. Quantification of mutant R150X expression in presence and absence of proteasomal inhibitor MG132
